# Most L1CAM is not associated with extracellular vesicles in human biofluids and iPSC–derived neurons

**DOI:** 10.1101/2024.10.14.618132

**Authors:** Vaibhavi Kadam, Madeleine Wacker, Patrick Oeckl, Milena Korneck, Benjamin Dannenmann, Julia Skokowa, Stefan Hauser, Markus Otto, Matthis Synofzik, David Mengel

**Affiliations:** Department of Neurodegenerative Diseases, Hertie Institute for Clinical Brain Research and Center of Neurology, University of Tuebingen, Tuebingen, Germany; German Center for Neurodegenerative Diseases (DZNE) Tuebingen, Tuebingen, Germany; Graduate School of Cellular and Molecular Neuroscience, University of Tuebingen, Tuebingen, Germany; Department of Neurology, Ulm University Hospital, Ulm, Germany; German Center for Neurodegenerative Diseases (DZNE) Ulm, Ulm, Germany; Department of Oncology, Hematology, Immunology, and Rheumatology, University Hospital Tuebingen, Tuebingen, Germany; Department of Neurology, Martin-Luther-University Halle-Wittenberg, Halle (Saale), Germany

**Keywords:** extracellular vesicles, L1CAM, biomarkers, neuron, cerebrospinal fluid, blood, isolation methods, immunoprecipitation

## Abstract

Transmembrane L1 cell adhesion molecule (L1CAM) is widely used as a marker to enrich for neuron-derived extracellular vesicles (EVs), especially in plasma. However, this approach lacks sufficient robust validation. This study aimed to assess whether human biofluids are indeed enriched for EVs, particularly neuron-derived EVs, by L1CAM immunoaffinity, utilizing multiple sources (plasma, CSF, conditioned media from iPSC-derived neurons [iNCM]) and different methods (mass spectrometry [MS], nanoparticle tracking analysis [NTA]).

Following a systematic multi-step validation approach, we confirmed isolation of generic EV populations using size-exclusion chromatography (SEC) and polymer-aided precipitation (PPT) – two most commonly applied EV isolation methods – from all sources. Neurofilament light (NfL) was detected in both CSF and blood-derived EVs, indicating their neuronal origin. However, L1CAM immunoprecipitation did not yield enrichment of L1CAM in EV fractions. Instead, it was predominantly found in its free-floating form. Additionally, MS-based proteomic analysis of CSF-derived EVs also did not show L1CAM enrichment.

Our study validates EV isolation from diverse biofluid sources by several isolation approaches and confirms that some EV subpopulations in human biofluids are of neuronal origin. Thorough testing across multiple sources by different orthogonal methods, however, does not support L1CAM as a marker to reliably enrich for a specific subpopulation of EVs, particularly of neuronal origin.

## 1. Introduction

Extracellular vesicles (EVs) are cell-secreted membrane-bound nanoparticles that carry biomolecular cargo, including proteins, nucleic acids and lipids, and play critical roles in intercellular communication and cell signaling [1–4], particularly in the context of brain health and disease [5, 6]. Given their ability to reflect physiological and pathological states of otherwise relatively inaccessible tissues such as those in the central nervous system (CNS), EVs hold large potential as biomarkers for various diseases, including neurological disorders. EVs are secreted into biological fluids such as blood and cerebrospinal fluid (CSF) [7], rendering them a promising tool for low-invasive fluid biomarker discovery in neurological disorders [8–11]. Yet the ability to reliably isolate and characterize EVs - especially those derived from neurons - is critical for validly utilizing their biomarker potential in neurological diseases. The transmembrane L1 cell adhesion molecule (L1CAM also known as, NCAM-L1 or CD171) [12] is the most frequently used neuronal EV marker to isolate and characterize EVs of presumed neuronal origin from human biofluids, particularly blood, due to its abundant expression reported in neuronal cells [13]. L1CAM has been employed in a relatively straightforward, medium-throughput immunoprecipitation-aided approach to isolate neuronal EVs from human biofluids, particularly blood. This method has rapidly gained attention, leading to a large number of biomarker studies in neurodegenerative diseases which utilized L1CAM to assess neuronal markers in blood [14–23].

However, despite its widespread use, the efficacy and specificity of L1CAM as a marker for EVs – in particular of neuronal origin – has not yet been robustly validated [14]. Specifically, concerns have emerged regarding the reliability of L1CAM-based immunoaffinity capture methods for isolating EVs, as L1CAM may primarily exist in a cleaved form in CSF and blood, rather than being bound to EV membranes [24], raising the question whether it is a reliable marker for EVs at all. Furthermore, L1CAM is also expressed in other cell types, including oligodendrocytes and in particular also various non-CNS tissues [25], questioning its use as a marker presumably specific for neuronal EVs.

In light of these questions, our study aimed to thoroughly evaluate whether human biofluids are indeed enriched for EVs by L1CAM immunoaffinity - particularly for neuron-derived EVs. We devised a multi-step validation approach utilizing multiple sources (plasma, CSF, conditioned media from iPSC-derived neurons [iNCM]) and different, orthogonal methods (mass spectrometry [MS], nanoparticle tracking analysis [NTA]). Our results validate EV isolation from diverse biofluid sources, using two commonly applied EV isolation methods in parallel (size-exclusion chromatography [SEC]; polymer-aided precipitation [PPT]) as well as a comprehensive characterization approach that included quantification and sizing of EVs by NTA and evaluation of several EV and EV exclusion markers. Moreover, they confirm that at least some EV subpopulations in human biofluids are of neuronal origin, by detecting neurofilament light (NfL) – a well-established neuronal marker – within the isolated EVs. Although EV subpopulations of neuronal origin were thus present, we did not find a specific enrichment for L1CAM in the EV fraction, as demonstrated by different, orthogonal methods (MS, NTA). Our findings provide further evidence that L1CAM may not serve as a reliable marker to enrich for a specific subpopulation of EVs, particularly those of neuronal origin.

## 2. Materials and methods

### 2.1 Human induced pluripotent stem cell (iPSC) neurons (iN) culturing and maintenance

iPSCs were differentiated to neurons of cortical layers V and VI based on Shi et al, 2012 with minor modifications [26][27]. A schematic illustrating the experimental protocol is provided in **Fig. S1**. Briefly, iPSCs were seeded in 6-well plates coated with Matrigel, at a density of 3 x 10^5^ cells/cm^2^ in E8 medium supplemented with 10 µM Y-27632 (Selleckchem). To stimulate neural induction, medium was changed to 3N medium supplemented with 15 µM SB431542 (Sigma-Aldrich) and 500 nM LDN-193189 (Sigma-Aldrich) for 9 days, with medium replaced every day. From day after induction (DAI) 9 to DAI 11, the cells were further expanded with medium containing 20 ng/mL bFGF (Peprotech). From DAI 13 to DAI 15, 100 µg/mL heparin was added to the 3N medium. Until DAI 26, the 3N medium was changed every alternate day. On DAI 26, the cells were dissociated using Accutase (Thermo Fisher Scientific) and plated at a density of approximately 7 x 10^5^ cells/cm^2^, and on DAI 27 and 29, 3N medium was changed and supplemented with 10 µM PD0325901 (Tocris) and 10 µM DAPT (Sigma-Aldrich). From DAI 31, 3N medium was changed every alternate day. Neuronal marker expression was assessed by immunocytochemistry using CTIP2, TUJ and TBR1 and differentiation was considered successful if more than 80% of the neurons were CTIP2-positive. All cells were maintained at 37°C at 5% CO2.

### 2.2 Human embryoid body (EB)-based hematopoietic stem cells (iH) culturing and maintenance

Human iPSCs were cultured under feeder-free conditions on Geltrex™ LDEV-free, hESC-qualified, reduced growth factor basement membrane matrix (Thermo Fisher Scientific) coated plates in StemFlex medium (Thermo Fisher Scientific) supplemented with 1% penicillin/streptomycin. Cells were passaged every three to four days with a density of 2 x 10^5^ cells/mL. Healthy Donor (HD) C16 iPSC line was kindly provided by Prof. T. Moritz and Dr. N. Lachmann, MHH Hannover. HD iPSC line was characterized elsewhere [28]. HD hiPSCs were dissociated from Geltrex™-coated plates at 70-80% confluency using PBS/EDTA (0.02%) for 7 min. Embryoid body (EB) formation was induced on day one via centrifugation of 20,000 cells per EB in 96-well plates with conical bottom (Sarstedt) using STEMdiff™ APEL™2 serum-free differentiation medium (StemCell Technologies) supplemented with bFGF (10 ng/ml) (Peprotech) and ROCK Inhibitor Y-27632 dihydrochloride (10 µM) (Tocris). BMP4 (40 ng/ml) (BioLegend) was added 24 h after EB formation to the culture to induce mesodermal differentiation. On day 4, EBs were plated on Geltrex™-coated 6-well plates (10 EBs/well) in APEL™2 medium supplemented with VEGF (40 ng/ml) (BioLegend), SCF (50 ng/ml) (R&D) and IL-3 (50 ng/ml) (BioLegend). For neutrophilic differentiation, medium was changed on day 7 to fresh APEL medium supplemented with IL-3 (50 ng/ml) (BioLegend) and G-CSF (10 ng/ml) (Filgrastim Hexal). The first hematopoietic suspension cells appeared on days 8 - 10. Differentiation medium was refreshed every three to four days with APEL medium supplemented with IL-3 (50 ng/ml) and G-CSF (10 ng/ml) until day 28 of hiPSC differentiation. Suspension cells were harvested and analyzed by flow cytometry on day 14 (CD34^+^ cells) and day 28 (neutrophils) as described before [29].

### 2.3 Extracellular vesicle isolation from cell conditioned media (iN and iH)

All steps to process the conditioned medium were performed at 4°C. Conditioned media from iN (iNCM) between days 27-40 and for iH (iHCM) at day 21 was collected and sequentially centrifuged at 4°C at 250 x g for 5 min, 400 x g for 5 min and finally at 2,000 x g for 10 min to discard pellets containing loose cells, cell debris and apoptotic bodies. Next, EDTA (pH 8.0) to a final concentration of 5 mM was added to the supernatant and mixed by inversion, and then stored at -80°C until use. On experiment day, media was thawed at 4°C, pooled and centrifuged at 2,000 x g for 10 min to pellet and discard any residual apoptotic bodies, followed by filtration with a 0.22 µm Express® Plus membrane (Merck Millipore). The filtered supernatant was concentrated to 0.5 mL using 10,000 Da MWCO 15 mL Amicon® ultra centrifugal filters (Millipore Sigma) pre-blocked with Tween-80 (Sigma-Aldrich) to mitigate non-specific binding, at 3,220 x g for 1 h to concentrate it to 0.5 mL. This 0.5 mL concentrate was loaded onto a qEVoriginal legacy size-exclusion chromatography (SEC) column (Izon Science) to collect 0.5 mL fractions between fractions 6 and 11 inclusive (EV fraction) and between fractions 12 and 20 inclusive (non-EV protein fraction). These EV and non-EV protein fractions were pooled and concentrated using 10,000 Da MWCO 2 mL Amicon® ultra centrifugal filters (Millipore) pre-blocked with Tween-80 (Sigma-Aldrich) at 3,220 x g for 30 min at 4°C to bring the volume to 100 µL each (for EV characterization experiments) or 0.5 mL and 1.0 mL respectively (for subsequent L1CAM immunocapture procedure) with 1X protease and phosphatase inhibitor cocktail (Halt™, Thermo Fisher Scientific) in Dulbecco’s Phosphate Buffered Saline (PBS) (Sigma-Aldrich).

### 2.4 Human plasma and CSF collection and handling

Human plasma and CSF samples were obtained from healthy donors from the biobank facility at the Hertie Institute for Clinical Brain Research, Tuebingen, Germany, and 1 mL and 0.5 mL volumes, respectively, were pre-aliquoted into LoBind tubes (Eppendorf) and stored at -80°C. They were thawed at 4°C only prior to EV isolation. For differential ultracentrifugation, a pooled human CSF sample (25 mL) was obtained from the biobank of the Department of Neurology, Ulm University Hospital (approval no. 20/10).

### 2.5 Extracellular vesicle isolation from CSF

CSF samples were thawed at 4°C and 5.5 mL was pooled per experiment, out of which 0.5mL was separately stored at -80°C for unprocessed CSF analyses. The remaining 5mL pooled CSF was subjected to centrifugation at 2,000 x g for 10 min at 4°C to remove cell debris and apoptotic bodies, after which the obtained supernatant was either subjected to EV isolation by SEC or precipitation using polyethylene glycol (PPT).

For EV isolation by PPT, 5 mL supernatant processed as above was gently mixed with 1 mL of ExoQuick (System Bioscience) each and incubated quiescent at 4°C overnight and then centrifuged at 1,500 x g for 20 min at 4°C to retrieve EVs in the pellet, which were resuspended and vortexed in 500 µL DPBS with 1.5X protease and phosphatase inhibitor cocktail, followed by facilitating overnight solubilization on a static rotator at 4°C. On the other hand, the supernatant was concentrated using 10,000 Da MWCO filters pre-blocked with Tween-80 at 3220 x g for 30 min at 4°C to bring the volume to 100 µL with 1.5X protease and phosphatase inhibitor cocktail in PBS.

For EV isolation by SEC, 5 mL supernatant was concentrated to 0.5 mL using 10,000 Da MWCO filters (Millipore Sigma) pre-blocked with Tween-80 (Sigma-Aldrich) at 3,220 x g for 30 min at 4°C. This concentrate was then loaded on a qEVoriginal legacy (Izon Science) column to collect 0.5mL fractions between fractions 6 and 11 inclusive (EV fraction) and between fractions 12 and 20 inclusive (non-EV protein fraction). The EV and non-EV protein fractions were pooled and concentrated using 10,000 Da MWCO filters (Millipore Sigma) pre-blocked with Tween-80 (Sigma-Aldrich) at 3,220 x g for 30 min at 4°C to bring the volume to 100 µL each with 1X protease and phosphatase inhibitor cocktail (Halt™, Thermo Fisher Scientific) in PBS (Sigma-Aldrich).

### 2.6 Extracellular vesicle isolation from CSF using differential ultracentrifugation (UC)

One pooled CSF sample (25 mL) underwent serial centrifugation to remove cells, debris and apoptotic bodies at 300 x g for 10 min, 2,000 x g for 10 min and 10,000 x g for 30 min, all at 4°C. An aliquot (1 mL) of the cleared CSF was aliquoted and stored for proteomic measurements (input CSF) and the remaining CSF volume was concentrated by 50 kDa Amicon centrifugal filters (Millipore). EVs were pelleted by UC at 100,000 x g for 90 min at 4°C. Pellets were washed with PBS and centrifuged again at 100,000 x g for 60 min. The EV pellet was resuspended in 50 µL PBS and stored at -80°C until analysis.

### 2.7 Mass-spectrometry based proteomics of EVs derived from CSF by UC

CSF EVs were disrupted by sonication and all samples (input CSF and CSF EVs) were reduced and alkylated using TCEP (5 mM final) and iodacetamide (10 mM final) and buffer exchanged with 50 mM triethyl ammonium bicarbonate (TEAB). Trypsin/LysC (Promega) was added in a ratio of 50:1 (protein-to-enzyme) and digested overnight at 37°C. Digestion was stopped by addition of TFA and equal amounts of protein were loaded on the nanoLC-MS system. Samples were analyzed in triplicates. Peptides were separated using an UltiMate 3000 RSLCnano system and a PepMap100 C18, 20 × 0.075 mm, 3 μm trap column (Thermo) and a PepMap100 C18, 50 × 0.075 mm, 2 μm analytical column (Thermo). Peptides were infused into a QExactive mass spectrometer and measured with data-dependent acquisition (Top12). Data were analyzed using MaxQuant v1.5.2.8 [30] and Perseus v1.5.2.6 [31]. N-terminal acetylation and methionine oxidation were set as variable modifications and carbamidomethylation on cysteine residues as fixed modification. Trypsin without cleavage before proline was set as the enzyme allowing up to two missed cleavages. Peptides and proteins were identified applying an FDR of 1%. For quantification, the MaxLFQ algorithm [32] was used. LFQ intensities were log2 transformed and missing values were replaced by imputation from a normal distribution (width 0.3, down shift 1.8). Groups were compared by Student’s t-test (two-tailed) and visualized by volcano plotting. Correction for multiple testing was performed with permutation-based FDR (0.05).

### 2.8 Extracellular vesicle isolation from plasma

Briefly, plasma samples were thawed at 4°C. 5 µL of thrombin (System Bioscience) was gently mixed with 500 µL plasma aliquot each and incubated at RT for 30 min, to remove fibrinogen and other clotting proteins, followed by addition of 500 µL PBS (Sigma-Aldrich) with 3X protease and phosphatase inhibitor cocktail (Halt™, Thermo Fisher Scientific). The reaction mixture was then centrifuged at 6,000 x g for 20 min at 4°C. The supernatants thus obtained were pooled and transferred to a fresh tube, and EVs were isolated by either SEC or PPT.

For EV isolation by SEC, the pooled supernatant was concentrated using 10,000 Da MWCO filters pre-blocked with Tween-80 at 3220 x g for 30 min at 4°C to bring the volume to 500 µL with 1.5X protease and phosphatase inhibitor cocktail in PBS and loaded on a qEVoriginal legacy (Izon Science) column to collect 0.5 mL fractions between fractions 6 and 11 inclusive (EV fraction) and between fractions 12 and 20 inclusive (non-EV protein fraction). These EV and non-EV protein fractions were pooled and concentrated using 10,000 Da MWCO filters pre-blocked with Tween-80 at 3,220 x g for 30 min at 4°C to bring the volume to 0.5 mL and 1 mL respectively with 1.5X protease and phosphatase inhibitor cocktail in PBS.

For EV isolation by PPT, 500 µL supernatant was gently mixed with 252 µL of ExoQuick (System Bioscience) each and incubated quiescent for 1 h at 4°C and then centrifuged at 1500 x g for 20 min at 4°C to retrieve EVs in the pellet which were resuspended and vortexed in 500 µL DPBS with 1.5X protease and phosphatase inhibitor cocktail, followed by facilitating overnight solubilization on a static rotator at 4°C. On the other hand, the supernatant was concentrated using 10,000 Da MWCO filters pre-blocked with Tween-80 at 3220 x g for 30 min at 4°C to bring the volume to 1 mL with 1.5X protease and phosphatase inhibitor cocktail in PBS.

### 2.9 NfL SIMOA measurements

Neurofilament light (NfL) concentrations were measured in singlet or technical duplicates on an ultrasensitive Single Molecule Array (SIMOA) on a Simoa HD-X analyzer (Quanterix), with NF-light Advantage kit (Quanterix, Lot#503729) used as per manufacturer’s instructions. Human CSF, plasma and iNCM samples were spun at 10,000 x g for 5 min at 4°C and then diluted using NfL sample buffer (Quanterix) for measurement. The lower limit of quantitation (LLoQ) of the assay was defined as the lowest standard (1) with a signal higher than the average signal for the blank plus 9 standard deviations, and (2) that allows a percentage recovery ≥100±20% and was determined to be 0.398 pg/mL for all measurements.

The NfL assay repeatability (%CV) for two internal control blood samples was determined as 3.3% and 6.4%.

### 2.10 L1CAM immunocapture

The method to enrich for L1CAM-associated EVs from plasma was adapted from previously established protocols [33]. 0.5 mL of EV fraction (corresponding to 0.5 mL plasma or 25 mL iNCM was resuspended with either 4 µg biotinylated anti-L1CAM antibody (Clone eBio5G3, Cat# 13-1719-82, eBioscience™, Invitrogen) or a home-made biotinylated anti-Calnexin antibody (Cat# ADI-SPA-860, Enzo Life Sciences) as a negative control in a total of 50 µL 3% bovine serum albumin in PBS (BSA/PBS). As a control for capturing free-floating L1CAM, 1 mL non-EV protein fraction (corresponding to 0.5 mL plasma or 25 mL iNCM) was resuspended in 8 µg biotinylated anti-L1CAM antibody or biotinylated anti-Calnexin antibody in a total 100 µL 3% BSA/PBS. All samples were mixed on a static rotator overnight at 4°C.

Next day, biotinylated EV and non-EV protein samples was incubated with 15 µL or 30 µL of pre-washed Streptavidin-Plus UltraLink Resin (Cat# 53116, Thermo Fisher) in 25 µL or 50 µL of 3% BSA/PBS for 4 h on a static rotator at 4°C. The resulting pellet containing L1CAM bound to the biotinylated antibody-resin complex was procured by two centrifugation steps at 400 x g for 10 min at 4°C, the second one being a wash step with 3% BSA/PBS. This was followed by elution with 100 µL 0.1 M glycine (pH 3.0) and an immediate centrifugation step at 4,500 x g for 5 min at 4°C to carefully retrieve the supernatant. Finally, 7.5 µL of 1 M Tris HCl (pH 8.0) in 12.5 µL 3% BSA/PBS was added to neutralize the suspension.

### 2.11 Nanoparticle tracking analysis (NTA)

Measurements were performed on NanoSight NS300 (Malvern Panalytical) NTA version 3.4 Build 3.4.003 to determine particle size and concentration. EVs were freshly prepared and maintained at 4°C and measured immediately. All samples were diluted with PBS. EVs from plasma samples were diluted 1/2000 while the immunoprecipitated samples were diluted 1/20. EVs from iNCM and CSF were diluted 1/500 and 1/50 respectively. Samples were measured in 5-10 technical replicates of 60 s each at camera level 15 and analyzed at detection threshold value 3. Particle count bin size was 5 nm, and particle concentration and size distribution profiles were plotted by intra-sample averaging of the replicates.

### 2.12 Protein concentration measurement and immunoblotting

Samples were treated with 1X RIPA buffer (150 mM NaCl, 1.0% IGEPAL® CA-630, 0.5% sodium deoxycholate, 0.1% SDS, 50 mM Tris, pH 8.0) and measured using the Pierce™ Bicinchonic Protein assay (ThermoFisher). Samples were denatured with 10X NuPAGE™ Reducing agent (Invitrogen) and 4X LDS sample buffer (Invitrogen) added to a final concentration of 1X each and subjected to heating at 70°C for 10 min. Proteins were separated by electrophoresis on NuPAGE™ Bis-Tris 10% 1.5 mm thickness mini or 4-12% 1.0 mm thickness midi gels at a constant voltage of 150 V and transferred at 4°C for 18 h at 0.05A or at RT for 30 min at 25 V onto 0.45µm pore size PVDF membranes. 100 µg of proteins from hIPSc-neuronal cell lysates, mouse brain lysates as well as whole plasma or CSF were loaded as positive controls. For western blotting involving plasma EVs, 37.5% of the volume equivalent of 0.5 mL plasma was loaded onto the gel to allow equivalent loading and direct comparison of IP-ed fractions and the pre-IP EV control. For western blotting involving iNCM-derived EVs, 12mL equivalent was loaded across all samples to be compared. TotalStain Q (Azure biosystems) total protein stain was used as a loading control and used as per manufacturer’s instructions. Membranes were blocked in Intercept™ blocking buffer-TBS (Li-COR Biosciences) and incubated with primary antibodies (ref. **Table 1** for detailed information on antibodies used) for L1CAM (clone UJ127, Cat# MA5-14140, Invitrogen), CD81 (clone B-11, Cat# sc-166029, Santa Cruz), Calnexin (Cat# ADI-SPA-860, Enzo Life Sciences), TSG101 (Clone 51/TSG101, Cat# 612696, BD Biosciences), Flotillin-1 (Clone 18/Flotillin-1, Cat# 610820, BD Biosciences), GM130 (Clone 35/GM130, 610823, BD Biosciences), either overnight at 4°C or at RT for 2 h in 1:1 Intercept™ blocking buffer and Tris-buffered saline with 0.1% Tween-20 (TBST), followed by the appropriate IRDye secondary antibodies at 1:10,000 for 1 h at RT, and visualized using the Li-COR Odyssey CLx imaging system (Li-COR Biosciences).

### 2.13 Data visualization

Data was visualized using GraphPad Prism v10.

## 3. Results

### 3.1 Size-exclusion chromatography (SEC) allows reliable isolation of EVs from iPSC-derived neurons and human biofluids

Due to its high specificity, recovery and scalability to isolate biophysically intact EVs, SEC is increasingly becoming the most convenient and practicable technique for biomarker research [4, 34]. As human biofluids contain heterogeneous populations of EVs secreted from various cell types, we first optimized our EV isolation procedure using SEC in a simplified model system. We used media conditioned by mature iPSC-derived neuronal cells (iNCM), a *bona fide* source of EVs secreted from neurons (**Fig. S1)**. Isolation of EVs from iNCM yielded particles in the size range of 40-200 nm as measured by nanoparticle tracking analysis (NTA), representing typical known size profiles of both exosomes as well as ectosomes [3] (**Fig. 1a**). Western blotting analysis confirmed presence of well-validated [35] EV markers CD81 and Flotillin-1, while the EV exclusion markers Calnexin and GM130 were absent (**Fig. 1a**), thus confirming the purity of the isolated EV populations. Following validation of the isolation protocol in EVs from iNCM, it was next applied to isolate EVs also from human CSF and blood. EVs isolated from both CSF and plasma using our protocol validated in iPSC-derived neurons, showed similar characteristics and presence of EVs markers, thus confirming that SEC reliably allows for the isolation of EVs from human biofluids (**Fig. 1b,c**).

**Fig. 1.**
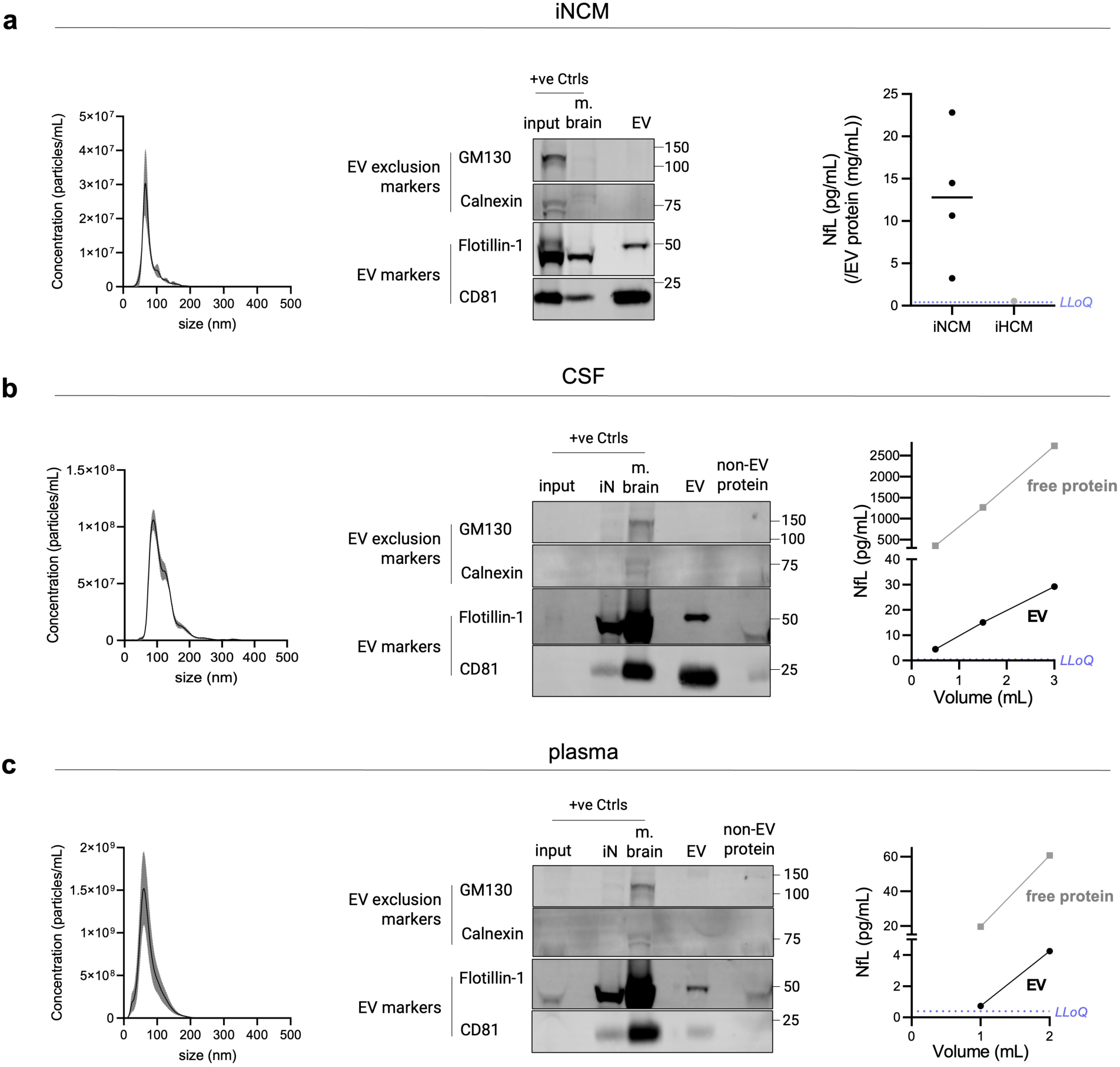
Isolation of EVs using size-exclusion chromatography (SEC) from conditioned medium of human induced pluripotent stem cells (iPSC-) neurons (iNCM) and biofluids consistently yields vesicles showing typical EV characteristics. EVs were isolated using SEC from **(a)** iNCM **(b)** CSF and **(c)** plasma and subjected to phenotypic characterization to confirm the presence of EVs. (**left**) Size distribution profiles of EV particles isolated from iNCM (n=3), plasma (n=5), CSF (n=3) using SEC measured by nanoparticle tracking analysis (NTA). Data is expressed as mean ± SEM. (**middle**) Representative immunoblots show the presence of classical EV markers (CD81, Flotillin-1) and absence of EV exclusion markers (Calnexin, GM130) in the EV fraction. Equal volumes of the non-EV protein fractions (to their respective EV fractions) were loaded to show absence of EV markers in them. For biofluids, positive controls include whole mouse brain lysate (m. brain) and iPSC-neuronal cell lysate (iN) and the unprocessed biofluid processed for EV isolation (input). For iNCM, whole mouse brain lysate (m. brain) and parent iPSC-neuronal lysate (input) were loaded as positive controls. Due to high sample viscosity, the conditioned medium and non-EV protein fraction could not be analyzed by immunoblotting. For each source, immunoblotting experiments were performed at least three times with similar results. (**right**) Neurofilament light chain (NfL) protein can be detected in iNCM, CSF and plasma, by SIMOA measurements. NfL can be detected in iNCM-derived EV NfL concentration (pg/mL) normalized to EV protein concentration (mg/mL), with each data point (black) representing a different iN cell line (n=4). EVs derived from conditioned medium of iPSC-derived hematopoietic stem cells (iHCM, grey) were used as a negative control for the setup (n=1) and show a much lower NfL signal than iNCM EVs. Due to high sample viscosity and interference of phenol-red present, the non-EV protein fraction of iNCM could not be measured. NfL levels in CSF and plasma also increase with increasing starting volume of the biofluid. The line plots show concentrations (pg/mL) of EV-associated (black) and free-floating (grey) NfL vs their starting volume, from one experiment. In all plots, the purple dashed line represents the lower limit of quantitation (LLoQ) value

### 3.2 The neuronal protein NfL is present in EVs isolated from human CSF and blood

To demonstrate that NfL can be detected in EVs derived from neuronal cells and thereby serve as a marker of neuronal origin in EVs in human biofluids, we once again, used EVs isolated from iNCM, serving as a *bona fide* experimental source of purely human neuronal EVs. We could detect NfL in EVs secreted into conditioned media in a concentration of 3.24-22.81 pg/mL (normalized to EV protein content). EVs isolated

from conditioned media of iPSC-derived hematopoietic stem and progenitor cells (iHCM) were used as EVs of non-neuronal origin and processed in the same manner. As expected, only minor amounts of NfL were detected in EVs from these non-neuronal cells (0.55 pg/mL when normalized to EV protein content). Next, we assessed EVs from human biofluids for the presence of NfL. Following separation of EVs from CSF and blood using our SEC-based protocol, we could confirm the presence of NfL in the non-EV protein fraction of CSF (488.51 pg/mL per mL CSF) and plasma (19.54 pg/mL per mL plasma), in line with expected concentrations of free-floating NfL in human CSF and blood reported in previous studies [36–39]. NfL was detected in the EV fractions of both CSF and plasma (9.68 and 0.72 pg/mL per mL starting volume, respectively), albeit at much lower concentrations than in the non-EV protein fractions (**Fig. 1b,c**).

### 3.3 L1CAM immunocapture of SEC-isolated plasma EVs suggests predominant free-floating L1CAM outside of EVs

Having validated SEC as a reliable method to isolate EVs from human biofluids, including those containing NfL as an indicator of neuronal origin, we next isolated neuronal EVs from blood, which is the most easily accessible and common source for EV biomarker research, via immunoprecipitation of L1CAM as the most commonly used method [14]. For immunoprecipitation, we used an anti-L1CAM antibody specific to the N-terminal region of its ectodomain (clone 5G3), which is also the most commonly used antibody for L1CAM immunocapture [14] (for schematic overview of the workflow, see **Fig. 2a)**. The anti-Calnexin antibody served as a control, as Calnexin should not be present in EVs. NTA analysis revealed that the size distribution profiles of particles isolated from plasma by SEC and followed by IP with either L1CAM or Calnexin were highly similar. Thus, we could not observe a specific enrichment for EV fractions using the anti-L1CAM antibody (as a presumable EV marker) compared to Calnexin (as an EV exclusion marker) (**Fig. 2b**). Next, we conducted immunoprecipitation (IP) using both L1CAM and Calnexin antibodies on both SEC-isolated EV as well as non-EV protein fractions, followed by probing for L1CAM through western blotting analyses. As expected, the SEC EV-enriched fraction (prior to IP) showed the presence of EV markers and the absence of EV exclusion markers (**Fig. 2c**), confirming effective isolation of EVs. Following L1CAM-IP of the SEC-EV fraction, we found no enrichment of L1CAM (**Fig. 2c**). However, we detected L1CAM when performing L1CAM-IP on the EV-depleted plasma fraction (**Fig. 2c**). No enrichment for L1CAM after IP with the anti-Calnexin antibody from the EV-depleted plasma fraction confirmed the specificity of this finding (**Fig. 2c**). In summary, upon enriching for L1CAM via IP, we found that the majority of L1CAM in plasma is present outside of EVs in a free-floating form rather than being associated with EVs.

**Fig. 2.**
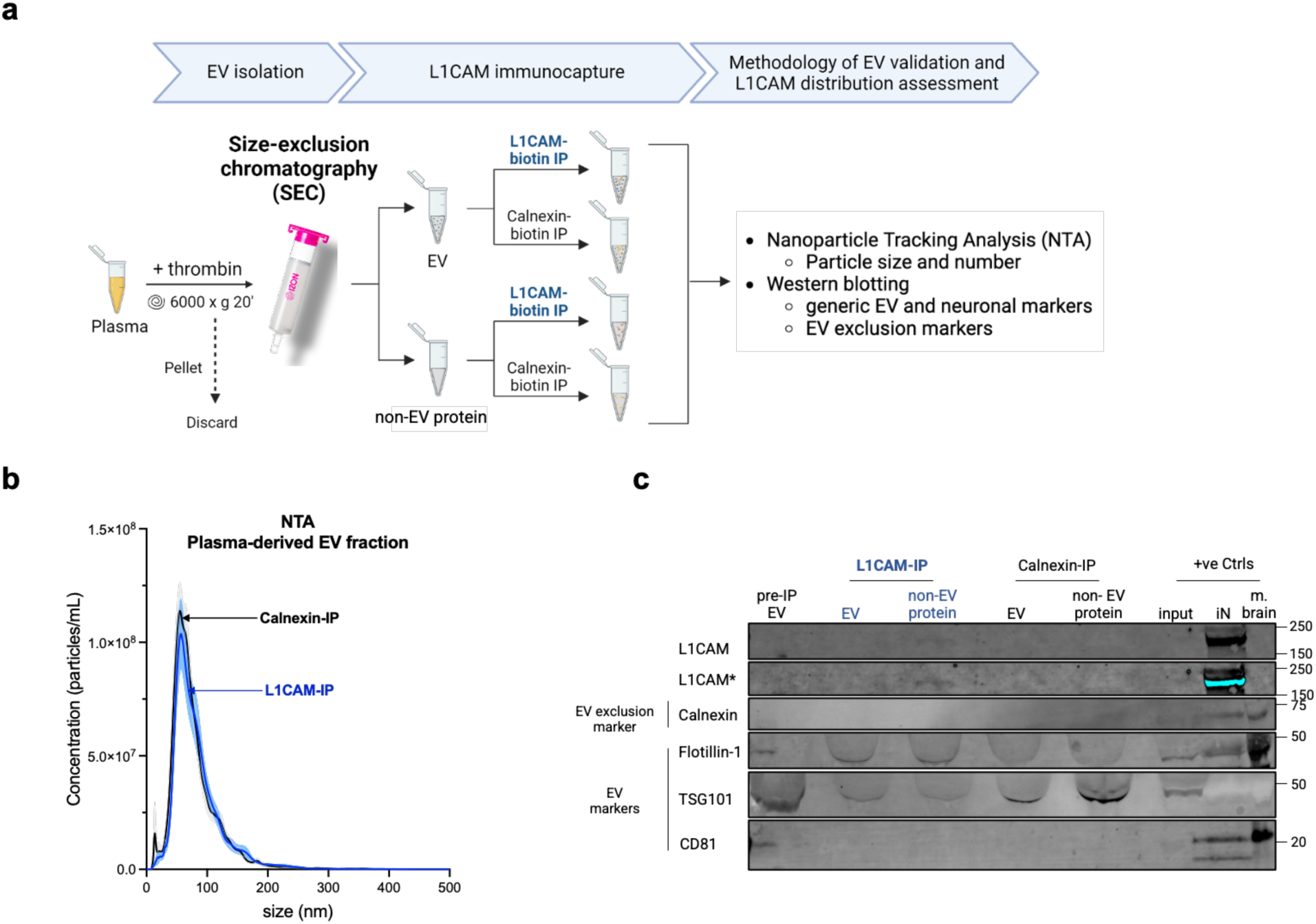
L1CAM is mostly found outside of EVs in its free-floating form, using size-exclusion chromatography (SEC)-based isolation and L1CAM-immunoaffinity purification. **(a)** Schematic of the methodological design implemented depicting that EV and non-EV protein fractions were isolated from thrombin-treated plasma by SEC and immunoprecipitated with biotin-conjugated L1CAM or Calnexin (used as a negative control). Image created using BioRender™. **(b)** Size distribution profiles of these plasma-derived EV fractions after IP by L1CAM (blue) or Calnexin (black) measured by NTA show no enrichment of a specific fraction of EVs by L1CAM. Data represents mean ± SEM from eight independent experiments, with x-axis indicating particle size in nanometer, while the y-axis representing particle numbers in 1 mL of plasma. **(c)** Representative immunoblot suggests that L1CAM is predominantly free-floating: Equivalent volumes of all IP-ed fractions and pre-IP EV fraction were loaded for direct comparison. The presence of EV markers (Flotillin-1, TSG101, CD81) in the pre-IP EV fraction confirms isolation of EVs and the absence of EV exclusion marker (Calnexin) serves as a control for EV purity. The blot reveals a notably higher signal intensity for L1CAM in the free protein fraction following L1CAM IP, in contrast to its counterpart in the EV fraction. Signal for L1CAM was also intentionally overexposed (marked with *) to improve visualization. Positive controls (+ve Ctrls) - whole mouse brain lysate (m. brain) and iPSC-neuronal cell lysate (iN) and the unprocessed plasma used for EV isolation (input) – were included to validate the experimental setup. The blot was repeated at least four times with consistent findings

### 3.4 EVs isolated by polymer-aided precipitation show similar characteristics and phenotype as SEC-isolated EVs

Polyethylene glycol (PEG)-based precipitation (PPT) is the most commonly used method for isolating plasma-derived EVs, specifically for subsequent L1CAM immunocapture, as this method is highly scalable and allows for higher protein recovery compared to SEC [3]. After isolation of EVs from plasma using PPT, we could also consistently isolate EVs within the same size range as observed using SEC-based isolation (**Fig. 3a**). Furthermore, the purity of the EV preparations was confirmed by the absence of EV exclusion markers and the presence of EV markers (**Fig. 3b**). This further validates PEG-aided precipitation, alongside SEC, as an effective method for isolating EVs from plasma. Additionally, mirroring the SEC data, we also confirmed the presence of NfL upon PPT-based isolation of EVs from plasma (0.17 pg/mL per mL plasma) (**Fig. 3c**) as well as from CSF (16.27 pg/mL per mL CSF) **(Fig. S2)**. Notably, using both methods of isolation (SEC, PPT), NfL signal positively correlated with increasing input volumes of both biofluids (**Fig. 1b,c, 3c, S2**).

**Fig. 3.**
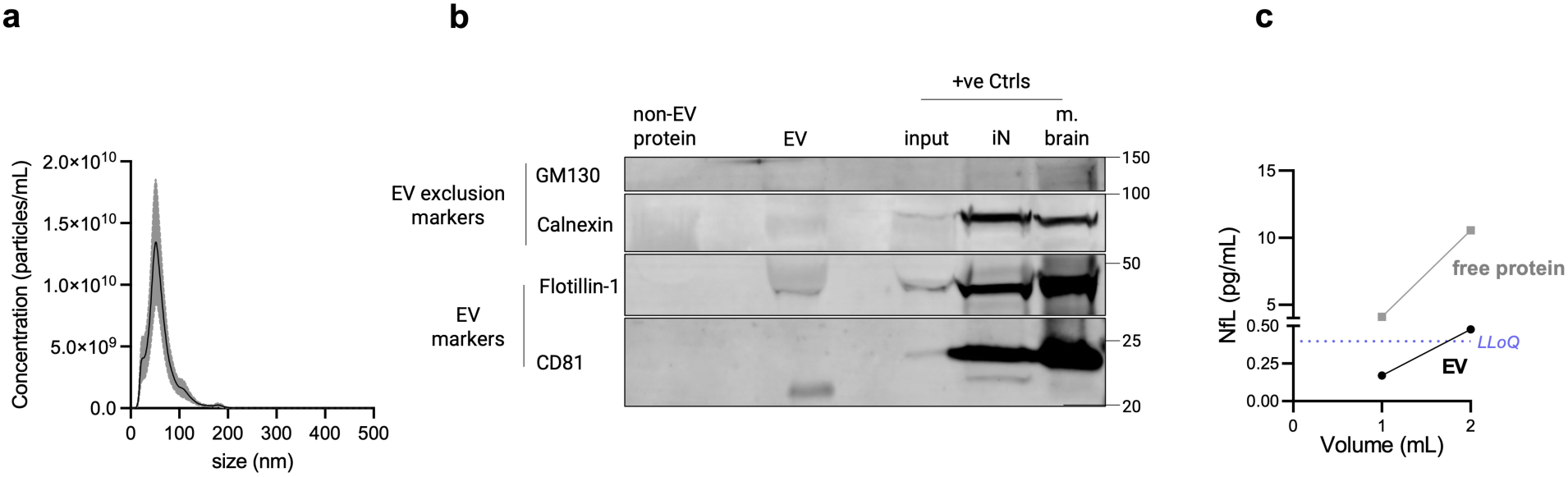
Isolation of EVs using polymer-aided precipitation (PPT) from plasma consistently yields EVs with typical characteristics, matching findings from SEC-based isolation. **(a)** Size distribution profiles of EV particles isolated from plasma measured by NTA. Data is expressed as mean ± SEM; n=6. **(b)** Representative immunoblot shows the presence of classical EV markers (CD81, Flotillin-1) and absence of EV exclusion markers (Calnexin, GM130) in the EV fraction. An equivalent volume of the non-EV protein fraction was also probed to check for absence of EV marker proteins. Whole mouse brain lysate (m. brain) and iPSC-neuronal cell lysate (iN) and unprocessed plasma used for EV isolation (input), were loaded as positive controls. The blot was repeated at least three times with similar results. **(c)** Neurofilament light chain (NfL) protein can be detected in plasma and increases with increasing starting volume. The line plots show concentrations (pg/mL) of EV- associated (black) and free-floating (grey) NfL as measured by SIMOA, with the purple dashed line denoting the lower limit of quantitation (LLoQ) value. This experiment was performed once

### 3.5 L1CAM is also found outside of EVs when immunoprecipitated from PPT-isolated EVs

Next, we investigated whether subsequent L1CAM immunocapture following PPT-based EV isolation could allow enrichment of an L1CAM-associated EV fraction, following a workflow (similar to SEC) as described in **Fig. 4a**. NTA analyses revealed similar size distribution profiles of particles immunoprecipitated by either anti-L1CAM or anti-Calnexin antibodies from PPT-isolated EVs (**Fig. 4b**), mirroring the results obtained with SEC-isolated EVs (**Fig. 2b**), and again suggesting no specific enrichment of certain fractions of EVs by L1CAM-IP. Immunoblotting analysis of PPT-isolated EVs and non-EV protein fractions followed by L1CAM-IP showed that L1CAM was mostly found outside of EVs (**Fig. 4c**), complementing our findings in SEC-isolated EVs (**Fig. 2c**).

**Fig. 4.**
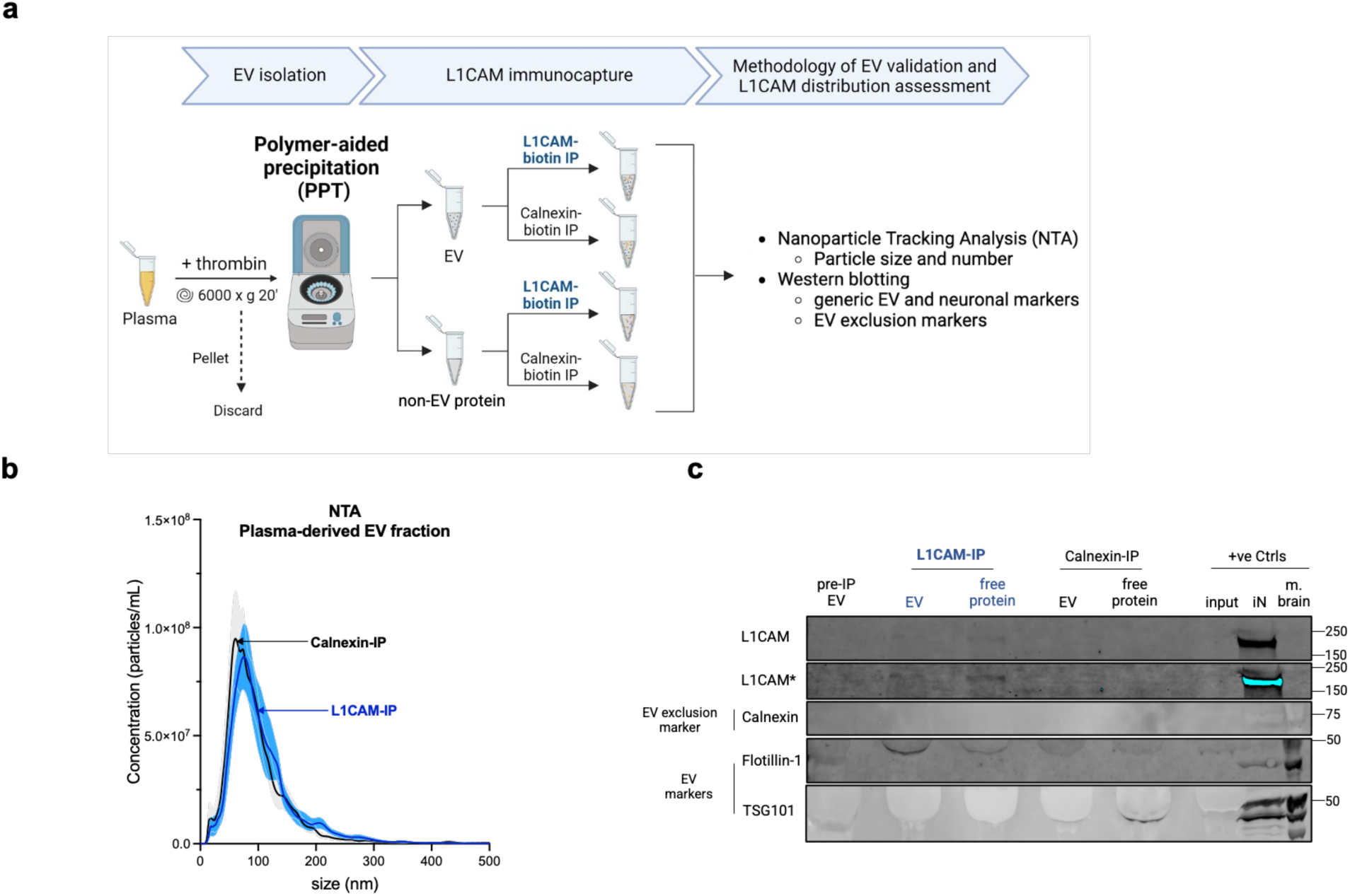
Most L1CAM is found free-floating compared to EV-associated in plasma, using the PPT isolation and immunocapture protocol mostly used in the field. **(a)** Schematic overview of the workflow showing that EV and non-EV protein fractions were first treated with thrombin and isolated from plasma by polymer-aided precipitation (PPT) and subsequently immunoprecipitated with biotin-conjugated L1CAM or Calnexin (used as a negative control). Image created using BioRender™. **(b)** Size distribution profiles of these plasma-derived EV fractions after IP by L1CAM (blue) or Calnexin (black) measured by NTA do not indicate enrichment of a specific fraction of EVs by L1CAM. Data represents mean ± SEM from seven independent experiments, with x-axis indicating particle size in nanometer, and the y-axis showing particle concentration (number of particles/mL) of input plasma. **(c)** Representative immunoblot suggests that most L1CAM is free-floating: Equivalent volumes of IP-ed fractions were loaded to allow direct comparison between them. Due to higher protein concentration of the pre-IP EV fraction, an approximately equivalent protein amount to IP-ed fractions was loaded to allow at least a qualitative degree of comparison. In the pre-IP EV fraction, the presence of EV markers (Flotillin-1, TSG101) confirms isolation of EVs and the absence of EV exclusion marker (Calnexin) controls for EV purity. L1CAM levels in the L1CAM-IPed non-EV protein fraction are higher than its EV fraction counterpart. Signal for L1CAM has also been shown overexposed (marked with *) to increase the signal to noise ratio and allow better visualization. Positive controls (+ve Ctrls) include whole mouse brain lysate (m. brain) and iPSC-neuronal cell lysate (iN) and unprocessed plasma used for EV isolation (input). Immunoblotting experiments were repeated at least four times and yielded similar results

Taken together, our data generated from plasma EVs isolated by two independent, but complementary methods (SEC and PPT) indicates that L1CAM is mostly present in its free-floating form outside of EVs and does not appear to be a suitable target to enrich for EVs (and thus, also not for neuron-derived EVs).

### 3.6 L1CAM-IP does not enrich for EVs present in the human iPSC-neuronal secretome

To further validate our findings also in a purely neuronal source of EVs, we next assessed the L1CAM- based IP protocol on EVs isolated from iNCM. Immunoblotting confirmed that the SEC-isolated EV fraction prior to IP showed the presence of EV markers (Flotillin-1, TSG101, CD81) and the absence of an EV exclusion marker (Calnexin) (**Fig. 5**). While L1CAM was present in the cell lysate of iPSC-derived neurons and showed some signal in their secreted EVs, this signal was absent in the EV fraction following L1CAM-IP - yet it was found in the non-EV protein fraction (**Fig. 5**). These results suggest that even in a purely neuronal source of EVs, there are higher levels of free-floating than EV-associated L1CAM, thus not supporting L1CAM-IP as a means to enrich for EVs of neuronal origin.

**Fig. 5.**
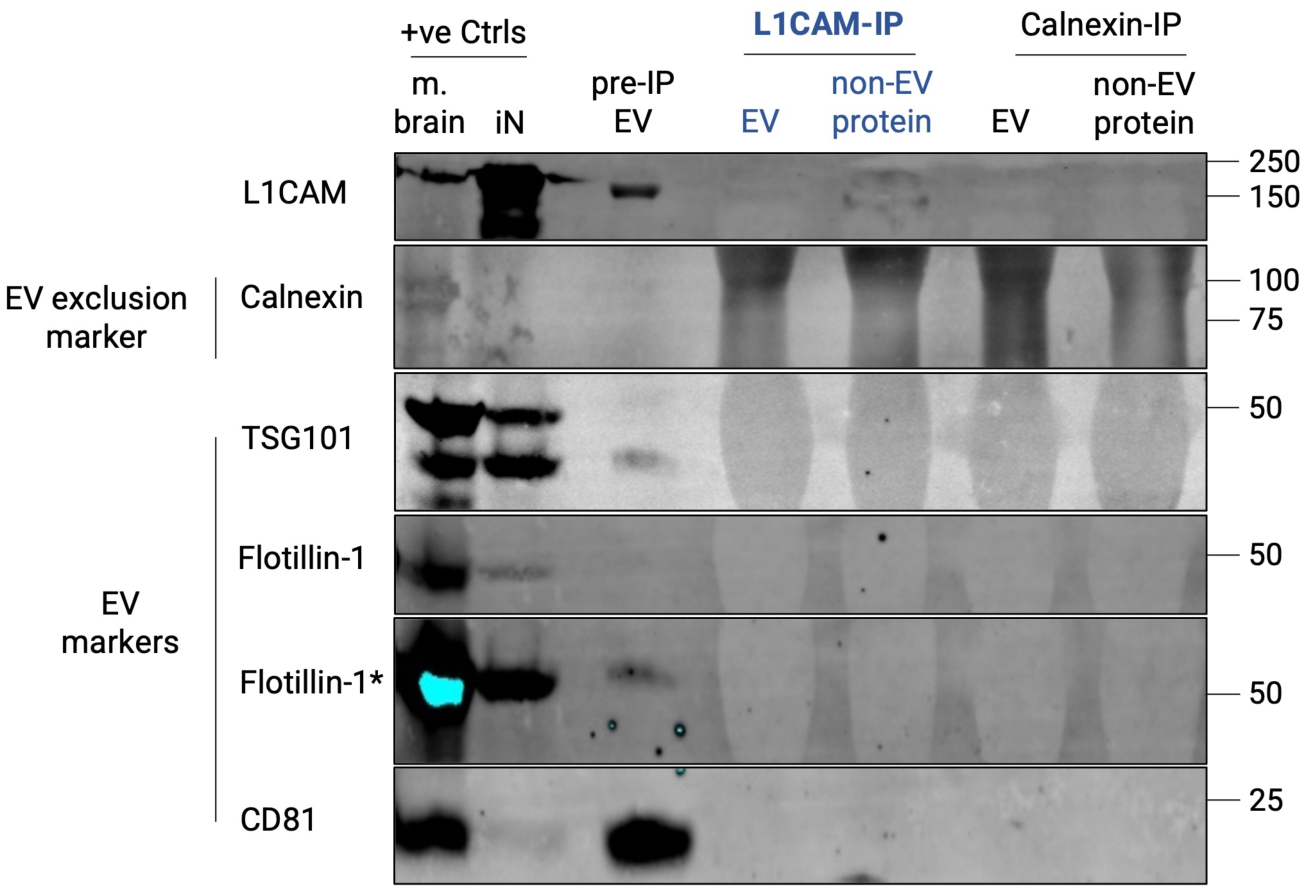
Higher levels of free L1CAM compared to EV-associated L1CAM after L1CAM immunoaffinity purification, in conditioned medium from iPSC-neurons. iPSC-neuronal conditioned medium was subjected to SEC-based isolation to separate EVs from other irrelevant and abundantly present proteins in the medium (e.g. albumin), before specifically enriching for L1CAM (L1CAM-IP) or Calnexin (used as a control). IP-ed EV and non-EV protein fractions as well as pre-IP EV (positive control) fractions were loaded by equivalent volume to allow their direct comparison. Effective isolation of EVs is confirmed by the presence of EV markers (Flotillin-1, TSG101, CD81) and the absence of EV exclusion marker (Calnexin) in the pre-IP EV fraction. The blot shows higher signal for L1CAM in the L1CAM-IPed non-EV protein fraction compared to its EV fraction counterpart. Positive controls (+ve Ctrls) - whole mouse brain lysate (m. brain) and iPSC-neuronal cell lysate (iN) were included to validate the experimental setup. Where required, the blot is overexposed to enable better visualization (marked as *). This experiment was repeated at least two times with similar results

### 3.7 L1CAM is not enriched in the proteome of EVs isolated from human CSF

Proteomic analyses present an alternative approach to assess the protein content of the EVs, complementing the analysis of particle size distribution by NTA and the detection of EV-enriched proteins by western blotting in the previous experiments. We hypothesized that L1CAM would be enriched in the proteome of EVs, alongside traditional EV markers, in EVs isolated from CSF, which is the biofluid closest to the brain, and hence most likely to contain neuronal-derived EVs. To allow best experimental conditions for proteomic analysis, EVs were harvested from CSF by differential centrifugation followed by ultracentrifugation (UC) – first at 300 x g, then 2000 x g, followed by 10,000 x g and lastly 100,000 x g [35]. Unbiased mass-spectrometry (MS)- based proteomics measurements performed on CSF-derived EVs showed high enrichment of canonical EV markers including CD9 (factor 207.94, *p* = 0.0033), CD81 (factor 225.97, *p* = 0.0043), Flotillin-1 (factor 28.25, *p* = 0.0016), TSG101 (factor 15.78, *p* = 0.0041) compared to unprocessed CSF (**Fig. 6**), indicating the validity of the approach to identify enrichment of EV markers. However, no specific enrichment of L1CAM was found in the EV fraction (factor 0.79, *p* = 0.71) (**Fig. 6**). These data provide additional evidence that L1CAM is not specifically associated with EVs and hence does not seem to be a suitable target for immunoprecipitation of EVs of anticipated neuronal origin.

**Fig. 6.**
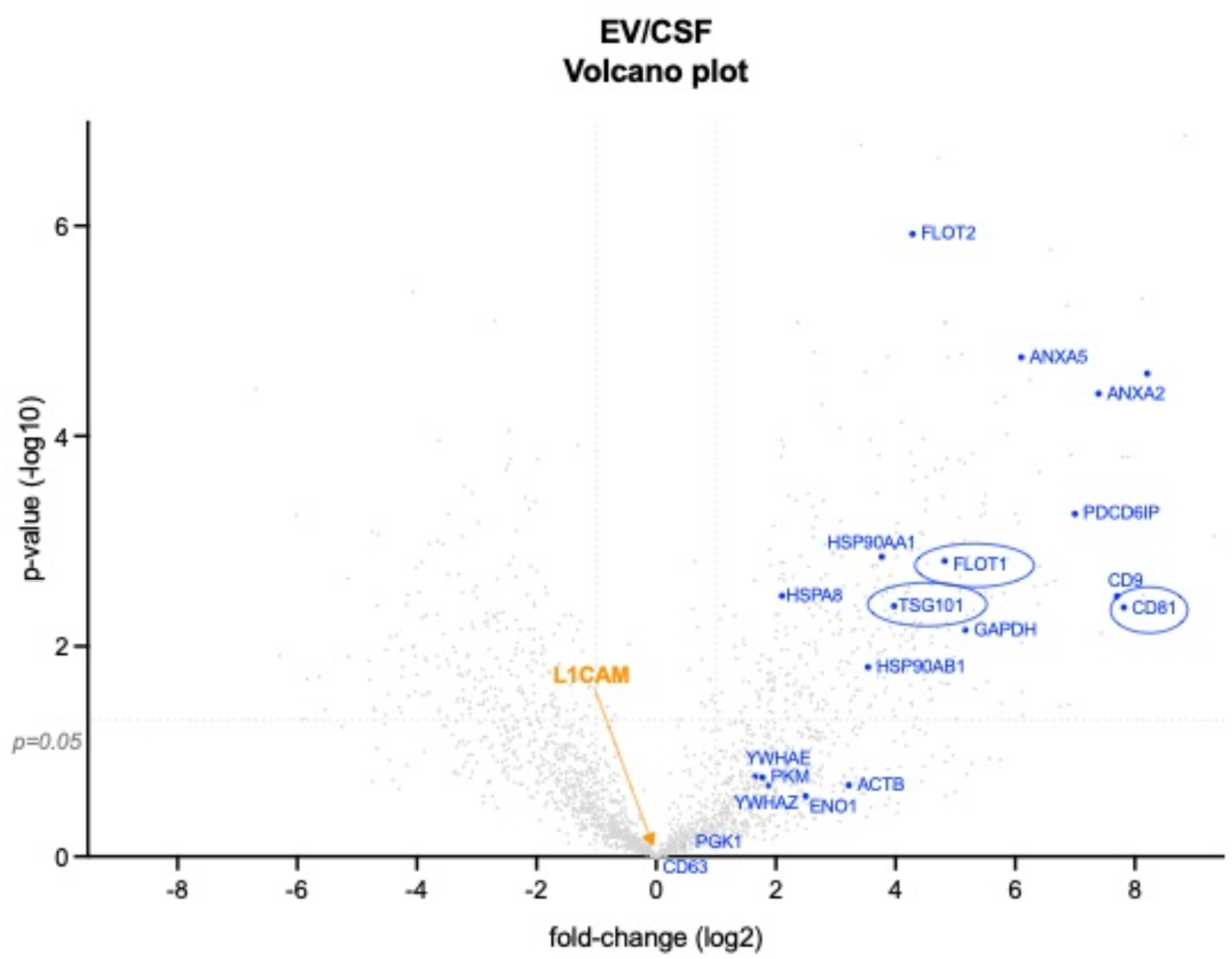
CSF-derived EVs isolated by differential ultracentrifugation (dUC) shows enrichment for canonical EV markers but not L1CAM. Volcano plot generated from unbiased mass-spectrometry (MS)-based proteomic results from CSF-derived EVs isolated by dUC, the current gold standard method for EV isolation, compared to unprocessed CSF, with x- and y-axes showing log2 transformed fold-change of EVs over CSF and –log10 transformed *p*-value, respectively, and the dashed solid lines depicting fold-change of 0.5 and 2 on the x-axis and *p* = 0.05 on the y-axis. The plot reveals specific enrichment of canonical EV markers (blue) with encircled ones used in the present study, while no enrichment of L1CAM (orange) is observed in the EV fraction. Data represents values from a single experiment

## 4. Discussion

The primary aim of this study was to confirm the *neuronal origin* of certain EV subpopulations within the broader EV populations derived from human biofluids; while, in addition, to critically test the reliability of L1CAM as a marker for enriching specific EV subpopulations - particularly those of *neuronal origin*.

### 4.1 Validation of SEC to enrich for generic EVs from human neurons, CSF, and blood

This study closely followed the guidelines outlined in the Minimal Information for Studies of Extracellular Vesicles (MISEV) 2024 [35] for EV nomenclature, storage, pre-processing, characterization and reporting. As part of a systematic stepwise validation approach, we first validated the isolation of *generic* EVs (= secreted from various tissues and cell types, and not specific to any cell type) from various human biofluids and culture media conditioned by cells. To achieve this, we initially validated a protocol for EV enrichment using SEC, as it has been previously identified as a reliable technique due to its superior specificity and recovery rates compared to other available methods [4, 34].

As a preliminary step, we isolated EVs from the conditioned media of iPSC-derived neurons (iNCM) to validate our method before applying it to more complex human biofluids, which contain a broad range of lipoproteins [40] and free proteins [41], that can complicate the reliable isolation of EVs. The EVs isolated from iNCM were confirmed to be within the expected size range (40-200 nm) for exosomes and ectosomes [3]. They contained well-established EV markers such as CD81 and Flotillin-1 [35], while EV exclusion markers like Calnexin and GM130 [35] were absent. Similarly, generic EVs enriched from the more complex human fluids- CSF and blood- met all quality criteria previously validated using EVs from cellular sources. Our findings align with those from other research groups that have isolated EVs from iNCM [42–44], CSF [45, 46], and plasma [47–50] using SEC, both in terms of EV sizes and EV markers. However, while CD81 and Flotillin-1 have been validated as reliable EV markers, many studies have not included EV exclusion markers (e.g., Calnexin, TOM20, GM130) to assess the purity of the EV preparation [48–50]. We here also incorporated such markers, as a critical quality-control measure to ensure that the detected signals are indeed from EVs - rather than from apoptotic bodies or cells.

### 4.2 Comparative isolation of EVs from blood using SEC and PPT

To demonstrate the generalizability of our findings beyond SEC-based EV isolation, we also applied a PPT-based EV isolation method- especially using plasma- as this method is most commonly used for isolating plasma-derived EVs, specifically for subsequent L1CAM immunocapture to enrich for neuron-derived EVs. This method is highly scalable, allows for higher protein recovery compared to SEC, and shows higher simplicity and cost-effectiveness, thus making it accessible for routine laboratory use. The isolated EVs consistently showed size profiles similar to our SEC-based results- typically ranging from 40-200 nm – which aligns with the expected size distribution for exosomes and other small vesicles. Purity was again confirmed by the presence of EV markers and the absence of exclusion markers. This validation across both SEC and PPT methods supports their reliability and efficiency in isolating EVs suitable for downstream applications, including biomarker discovery and therapeutic research.

### 4.3 NfL indicates presence of neuronally derived EVs in human CSF and blood

NfL is an intermediate filament protein specific to neurons, where it forms an essential component of the neuronal cytoskeleton [51–53]. As EVs in both CSF and blood can originate from a variety of tissues and cell types, we utilized expression of NfL as an indicator of those EV subpopulations that are derived from neuronal origin in both fluids. We detected NfL in EVs from iNCM, thereby confirming the sorting of NfL into EVs released from neurons and validating it as a marker of EVs of neuronal origin. The high expression of NfL in EVs from iNCM – with only minute amounts being present in EVs from iHCM - confirmed the specificity of this finding. The minimal but detectable signal for NfL in EVs from iHCM could be attributed to the early differentiation stage of these hematopoietic cells [29]. Additionally, bone marrow T-cells have also been suggested to have a basal NfL signal - albeit only analyzed at the RNA level [54].

We also detected NfL in EVs isolated from human CSF and blood, validating that a subpopulation even of generic EVs contained in human biofluids - and isolated by both methods- SEC and PPT- are indeed of neuronal origin. Although NfL levels were low in plasma-derived EVs (by both methods), its levels positively correlated with increasing input volume of plasma, also consistently demonstrated by both methods, suggesting that the observed signal is genuine and not merely due to e.g. only unspecific binding or background signal. The observed low levels of NfL are most likely just reflective of the inherently low concentrations of NfL within EV populations from peripheral sources, e.g. plasma samples, as opposed to CSF samples (as demonstrated here). Thus, in sum, by employing two different isolation methods - each widely used- we hereby systematically confirm and extend previous studies [45, 55–57], showing the presence of NfL in EVs from human biofluids.

### 4.4 L1CAM immunocapture of plasma EVs isolated by SEC and PPT suggests predominant free-floating L1CAM outside of EVs

Immunocapture of L1CAM, a prominent putative neuronal EV surface marker, has been extensively used in biomarker studies to enrich for neuron-derived EVs from human blood [14–23]. However, this approach lacks thorough methodological characterization and validation [14]. To this end, we first determined whether anti-L1CAM IP indeed enriches for L1CAM protein in a subpopulation of EVs following SEC/PPT-based EV isolation from blood, for which a successful isolation protocol had been established in the first step of our study. To control for unspecific enrichment by the IP procedure, we established an anti-Calnexin IP as a control condition, as Calnexin is an ER membrane protein and not sorted into EVs [35]. Following anti-L1CAM and anti-Calnexin IP in EV fractions of SEC and PPT, we could not find differences in the enrichment of certain populations of EVs by both methods, suggesting unspecific interaction between EVs and the immunocomplexes in the IP process. Furthermore, while L1CAM was not enriched by IP in EVs, it was detected in the corresponding non-EV protein fractions after IP, thus representing free-floating L1CAM. These data confirm and validate a previously published study that suggested that L1CAM is not associated with EVs in blood and CSF, but rather present in these fluids in its soluble form (e.g. due to cleavage and/or alternative splicing) [24]. Furthermore, we also extend these recent findings by employing immunocapture and enrichment techniques, - not only to enhance detection specificity for L1CAM but also to determine relative L1CAM levels (EV-associated vs free), again exactly following the standard procedure used for L1CAM-associated EV enrichment. While the absence of L1CAM in L1CAM-immunoprecipitated EVs does not exclude the possibility that minor subpopulations of L1CAM-positive EVs might still be present, it fails to provide positive evidence supporting this widely used approach involving PPT-based isolation followed by L1CAM immunocapture.

Nevertheless, L1CAM might still serve as a valid marker to enrich for EVs. In other words, while L1CAM immunocapture is expected to enrich for neuronal EVs amongst the predominantly peripheral EVs in blood, sensitivity may not be achieved owing to the L1CAM-positive EV subpopulation being too scarce in this CNS-distant compartment. We therefore hypothesized that enrichment of L1CAM may be higher in EVs directly released by a well-defined and purely neuronal source (e.g. iNCM), where a significant proportion of all EVs would be expected to express it, given the abundant neuronal cellular L1CAM expression. In line with our findings in plasma, EVs released in iNCM contained NfL (confirming their neuronal origin), but L1CAM was predominantly found free-floating after L1CAM IP and not associated with EVs. These findings add additional evidence that L1CAM is not abundantly present on EVs - here including even neuronal EVs that also contain NfL.

### 4.5 L1CAM is not enriched in the proteome of EVs isolated from human CSF

Proteomic analyses are a valuable approach to further assess the protein content of EVs, providing a more comprehensive understanding of their molecular composition. Our proteomic approach served as an independent, orthogonal analysis method to our previous analyses, which included particle size distribution via NTA and detection of EV-enriched proteins through western blotting. CSF is the biofluid most proximate to the brain and central nervous system, and therefore, it is considered the most likely source of neuron-derived EVs. Based on literature [58], we hypothesized that L1CAM would be enriched in the proteome of EVs isolated from CSF, alongside conventional EV markers such as CD81, Flotillin-1, TSG101. However, proteomic analysis revealed that CSF-derived EVs showed no significant enrichment for L1CAM compared to unprocessed CSF, while established EV markers (CD9, TSG101, CD63, Flotillin- 1, etc.) were highly enriched in CSF-derived EVs vs unprocessed. This is consistent with another proteomic study in human CSF EVs isolated by UC at 100,000 x g [59]. Our results also corroborate and extend a previous report which found no enrichment of L1CAM in canine CSF-derived EVs using SEC-based isolation, and majorly eluted with albumin in the non-vesicular fractions [46]. These findings add further evidence by an orthogonal method- that -unlike other EV markers- L1CAM is not enriched in EVs isolated from CSF.

### 4.6 Limitations

The results of our study need to be interpreted in light of several limitations. First, our findings on NfL in EVs isolated from CSF and blood suggest that a detectable proportion of EVs in these biofluids may be of neuronal origin. Future studies should expand on this by employing single particle analysis to investigate the colocalization of NfL with additional neuronal markers, such as MAP2 or synaptophysin using flow cytometry [60, 61] or immune-gold labelling electron microscopy [62]. These experiments - which were beyond the scope of this manuscript - could also be used to assess the colocalization of L1CAM with neuronal markers, providing further insight into whether a subset of neuronal EVs might contain surface L1CAM. Second, further research is warranted to explore L1CAM-positive EVs *in vivo* using transgenic mouse models that express L1CAM conjugated to a fluorophore. Biofluids, such as blood and CSF, extracted from these models could then be analyzed to determine the presence and relative proportion of L1CAM-positive vesicles compared to other EVs. Third, Norman and colleagues [24] have reported that the soluble L1CAM protein found in CSF and plasma may be cleaved or alternatively spliced or both. To build on this, future research should investigate the free-floating L1CAM proteoform that we consistently detect in our plasma and iNCM, blots. This could be achieved through mass-spectrometry-based proteomics to compare the peptide sequences of EV-associated vs free L1CAM forms in plasma, CSF, and their respective EVs to determine whether the soluble L1CAM is in its native full-length form or not.

### 4.7 Conclusions

This study systematically validated the isolation of generic EVs assessing various human biofluids, such as CSF, and blood as well as conditioned media from neurons- using two parallel and complementary isolation methods (SEC, PPT). We confirmed that these methods are effective for enriching EVs, as evidenced by the presence of established EV markers and the absence of EV exclusion markers. Our investigation into the neuronal origin of EV subpopulations revealed that NfL can serve as an indicator of neuron-derived EVs in both CSF and blood. However, our findings challenge the utility of L1CAM as a marker for EVs (and in particular also specifically as a marker for neuronal EVs), as L1CAM was not enriched in the EV fractions from these biofluids, but rather appeared predominantly in non-EV protein fractions. This suggests that L1CAM may not be as prevalent on neuronal EVs as previously assumed, prompting a need for further research into alternative EV markers – in particular of neuronal origin- and more nuanced isolation techniques.

## Acknowledgements

We thank Jacob Helm, Kalaivani Manibarathi, Linus Wiora and Maike Nagel (Hertie Institute for Clinical Brain Research, University of Tuebingen) for their generous help in providing media conditioned by human iPSC-neuronal cultures, and Stephen Meier (Department of Neurology, Ulm University Hospital) for his excellent technical assistance. We are also grateful to the Biobanks at the Hertie Institute for Clinical Brain Research, Tuebingen (Claudia Schulte, Christian Deuschle) and Department of Neurology, Ulm University Hospital (Alice Beer, Sandra Huebsch, Dagmar Schattauer) for providing blood and CSF samples.

## Conflicts of interest

Vaibhavi Kadam, Madeleine Wacker, Milena Korneck, Benjamin Dannenmann, Julia Skokowa, Stefan Hauser and David Mengel report no disclosures. Patrick Oeckl received research support from the Cure Alzheimer Fund, ALS Association (24-SGP-691, 23-PPG-674-2), ALS Finding a Cure, the Charcot Foundation, the DZNE Innovation-to-Application program and consulting fees from LifeArc and Fundamental Pharma, all unrelated to the present manuscript. Markus Otto reports served as scientific advisor for Biogen, Roche, Axon, Fuijrebio. He received research support from the Cure Alzheimer Fund, Boehringer Ingelheim University Ulm Institute, Thierry Latran foundation, European commission and the BMBF, all unrelated to the present manuscript. Matthis Synofzik has received consultancy honoraria from Ionis, UCB, Prevail, Orphazyme, Servier, Reata, GenOrph, AviadoBio, Biohaven, Solaxa, Biogen, Zevra, and Lilly, all unrelated to the present manuscript.

## Statements and Declarations Author contributions

Vaibhavi Kadam, David Mengel and Matthis Synofzik contributed to study conception and design. Material preparation, experimentation and analysis were performed by Vaibhavi Kadam, Madeleine Wacker, Patrick Oeckl, Milena Korneck and Benjamin Dannenmann. Vaibhavi Kadam, David Mengel and Matthis Synofzik wrote the initial draft of the manuscript. All authors contributed to writing the manuscript and read and approved the final manuscript.

## Funding

This work was supported by the Clinician Scientist program “PRECISE.net” funded by the Else Kröner-Fresenius-Stiftung (to D.M, and M.S.), by the EU Joint Programme - Neurodegenerative Disease Research GENFI-PROX grant [2019-02248; to M.S). D.M. is supported by the Clinician Scientist program of the Medical Faculty Tübingen (459-0-0) and the Elite Program for Postdoctoral researchers of the Baden-Württemberg-Foundation (1.16101.21).

## Informed consent statement

Informed consent was obtained from all subjects involved this study wherever necessary.

## Ethics approval

This study was performed in line with the principles of the Declaration of Helsinki and approved by the Ethics Committee of the University of Tuebingen (199/2011BO1).

## Data availability statement

All data supporting the findings of this study are available within this paper and its supplementary information. Further information can be provided upon reasonable request to the corresponding author.

## Notes

### Competing Interest Statement

The authors have declared no competing interest.

### Summary of Updates

The western blot panel of figure 4c has been revised

