## Supplementary information for "Most L1CAM is not associated with extracellular vesicles in human biofluids and iPSC–derived neurons"

**Supplementary figures**

**Fig. S1**

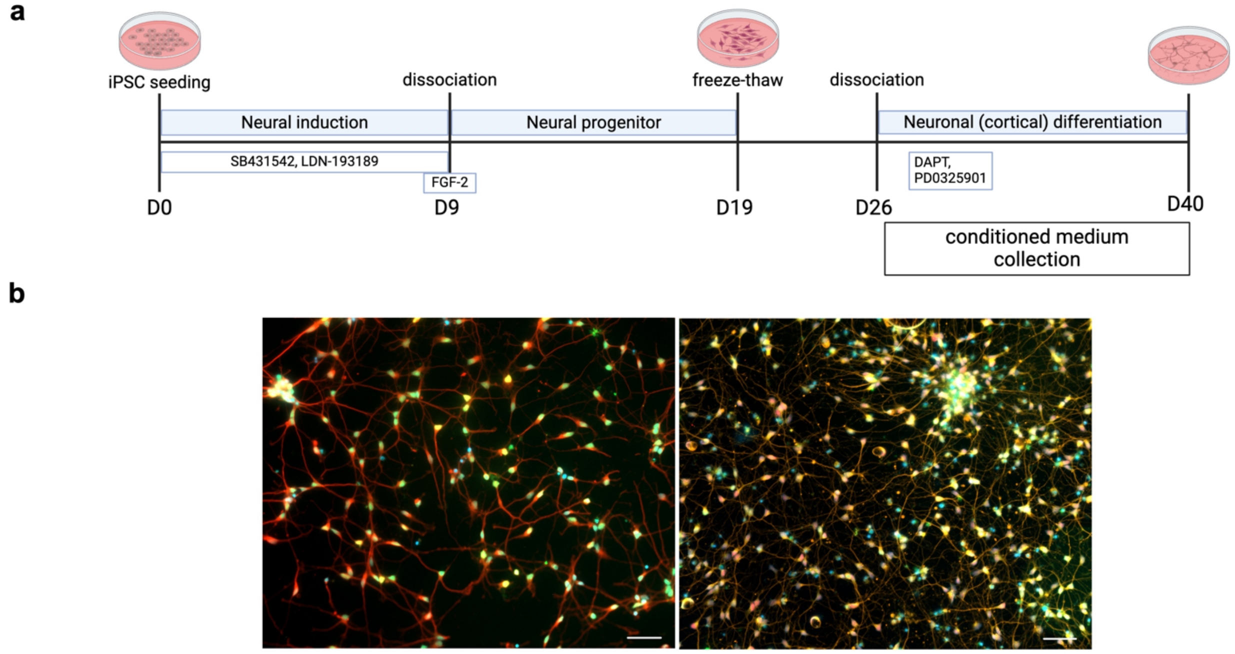


Fig. S1 **Immunocytochemistry (ICC) staining of human iPSC-derived cortical neurons shows presence of classical neuronal markers.**

**(a.)** Schematic diagram of the experimental protocol implemented for iPSC differentiation into iPSC-derived cortical neurons. Conditioned media for EV isolation was collected between DAI 27 and 40. **(b.)** Representative ICC images of iPSC-derived neurons stained for CTIP2 (layer V cortical neuron marker, in green), TBR1 (layer VI cortical neuron marker, in far-red), TUJ (neuronal differentiation marker, in red[left]/orange[right]) and DAPI (nuclear marker, in blue) at day 36 of neuronal differentiation. Magnification = 20X, scale bars = 50 μm

**Fig. S2**

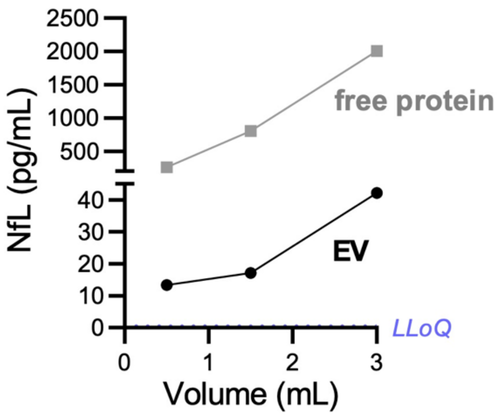


Fig. S2 **NfL levels in CSF-derived EVs isolated by PPT also increase with increasing CSF starting volume.**

The line plots denote SIMOA measurements of EV- associated (black) and free-floating (grey) NfL concentrations (pg/mL), with the purple dashed line represents the lower limit of quantitation (LLoQ) value (n=1)

**Tables**

**Table 1**

| Antibody | Host | Clonality (with clone) | Manufacturer | Dilution/ final concentration | Application |
|---|---|---|---|---|---|
| anti-L1CAM | mouse | Monoclonal (UJ127) | Invitrogen | 1:200 | WB |
| CD81 | mouse | Monoclonal (B-11) | Santa Cruz | 1:1000 | WB |
| Anti-Calnexin | rabbit | Polyclonal | Enzo Life Sciences | 1:2000 | WB |
| Anti-TSG101 | rabbit | Monoclonal (51/TSG101) | BD Biosciences | 1:1000 | WB |
| Anti-Flotillin-1 | mouse | Monoclonal (18/Flotillin-1) | BD Biosciences | 1:1000 | WB |
| Anti-GM130 | mouse | Monoclonal (35/GM130) | BD Biosciences | 1:1000 | WB |
| Anti-L1CAM (biotinylated) | mouse | Monoclonal (5G3) | Thermo Fisher Scientific | 16µg/mL | IP |
| Anti-Calnexin (biotinylated (home-made)) | rabbit | Polyclonal | Enzo Life Sciences | 16µg/mL | IP |

Table 1. **List of antibodies used.** *WB*, western blotting; *IP*, immunoprecipitation
